# Increased Resistance of SARS-CoV-2 Variant P.1 to Antibody Neutralization

**DOI:** 10.1101/2021.03.01.433466

**Authors:** Pengfei Wang, Ryan G. Casner, Manoj S. Nair, Maple Wang, Jian Yu, Gabriele Cerutti, Lihong Liu, Peter D. Kwong, Yaoxing Huang, Lawrence Shapiro, David D. Ho

## Abstract

The relative resistance of SARS-CoV-2 variants B.1.1.7 and B.1.351 to antibody neutralization has been described recently. We now report that another emergent variant from Brazil, P.1, is not only refractory to multiple neutralizing monoclonal antibodies, but also more resistant to neutralization by convalescent plasma (3.4 fold) and vaccinee sera (3.8-4.8 fold). The cryo-electron microscopy structure of a soluble prefusion-stabilized spike reveals the P.1 trimer to adopt exclusively a conformation in which one of the receptor-binding domains is in the “up” position, with the functional impact of mutations appearing to arise from local changes instead of global conformational alterations. The P.1 variant threatens current antibody therapies but less so the protective efficacy of our vaccines.

---

SARS-CoV-2 P.1, emerging from the B.1.1.28 lineage, has become a dominant variant in Brazil (Faria, 2021; Naveca, 2021). P.1 contains 10 spike mutations in addition to D614G, including K417T, E484K, and N501Y in the receptor-binding domain (RBD), L18F, T20N, P26S, D138Y and R190S in the N-terminal domain (NTD), and H655Y near the furin cleavage site. This new variant could threaten the efficacy of current monoclonal antibody (mAb) therapies or vaccines, because it shares mutations at the same three RBD residues with B.1.351, a variant that first emerged from South Africa (Tegally et al., 2021). We and others (Liu et al., 2021; Wang et al., 2021; Wu et al., 2021) have shown that B.1.351 is more resistant to neutralization by some mAbs, convalescent plasma, and vaccinee sera, in part due to a E484K mutation that also exists in P.1. We therefore obtained the P.1 authentic virus and also created, as previously described (Liu et al., 2020; Wang et al., 2020; Wang et al., 2021), a VSV-based SARS-CoV-2 pseudovirus with all 10 mutations of the P.1 variant (BZΔ10), and assessed their susceptibility to neutralization by 18 neutralizing mAbs, 20 convalescent plasma, and 22 vaccinee sera as previously reported (Wang et al., 2021).

We first assayed the neutralizing activity of four mAbs with emergency use authorization (EUA), including REGN10987 (imdevimab), REGN10933 (casirivimab) (Hansen et al., 2020), LY-CoV555 (bamlanivimab) (Chen et al., 2021; Gottlieb et al., 2021), and CB6 (etesevimab) (Gottlieb et al., 2021; Shi et al., 2020) against P.1 pseudovirus (BZΔ10) and authentic virus, alongside with their wildtype (WT or WA1) counterparts. As shown in Figure 1A (left panel) and Figure S1A, the neutralizing activities of three of the mAbs with EUA were markedly or completely abolished against P.1. The only mAb with EUA retaining its activity was REGN10987. We next tested the neutralizing activity of eight additional RBD mAbs, including ones from our own collection (2-15, 2-7, 1-57, & 2-36) (Liu et al., 2020) as well as S309 (Pinto et al., 2020), COV2-2196 & COV2-2130 (Zost et al., 2020), and C121 (Robbiani et al., 2020). The neutralizing activities of the two potent mAbs targeting the receptor-binding motif, 2-15 and C121, were completely lost against P.1 (Figure 1A, middle panel, and Figure S1A). Other mAbs targeting the “inner side” or the “outer side” of RBD retained their activities against P.1, however. Overall, the data on pseudovirus and authentic virus were in agreement, and the findings on P.1 mimic those observed for B.1.351 (Wang et al., 2021), which should not be surprising since the triple RBD mutations in P.1 and B.1.351 are largely the same.

**Figure 1.**
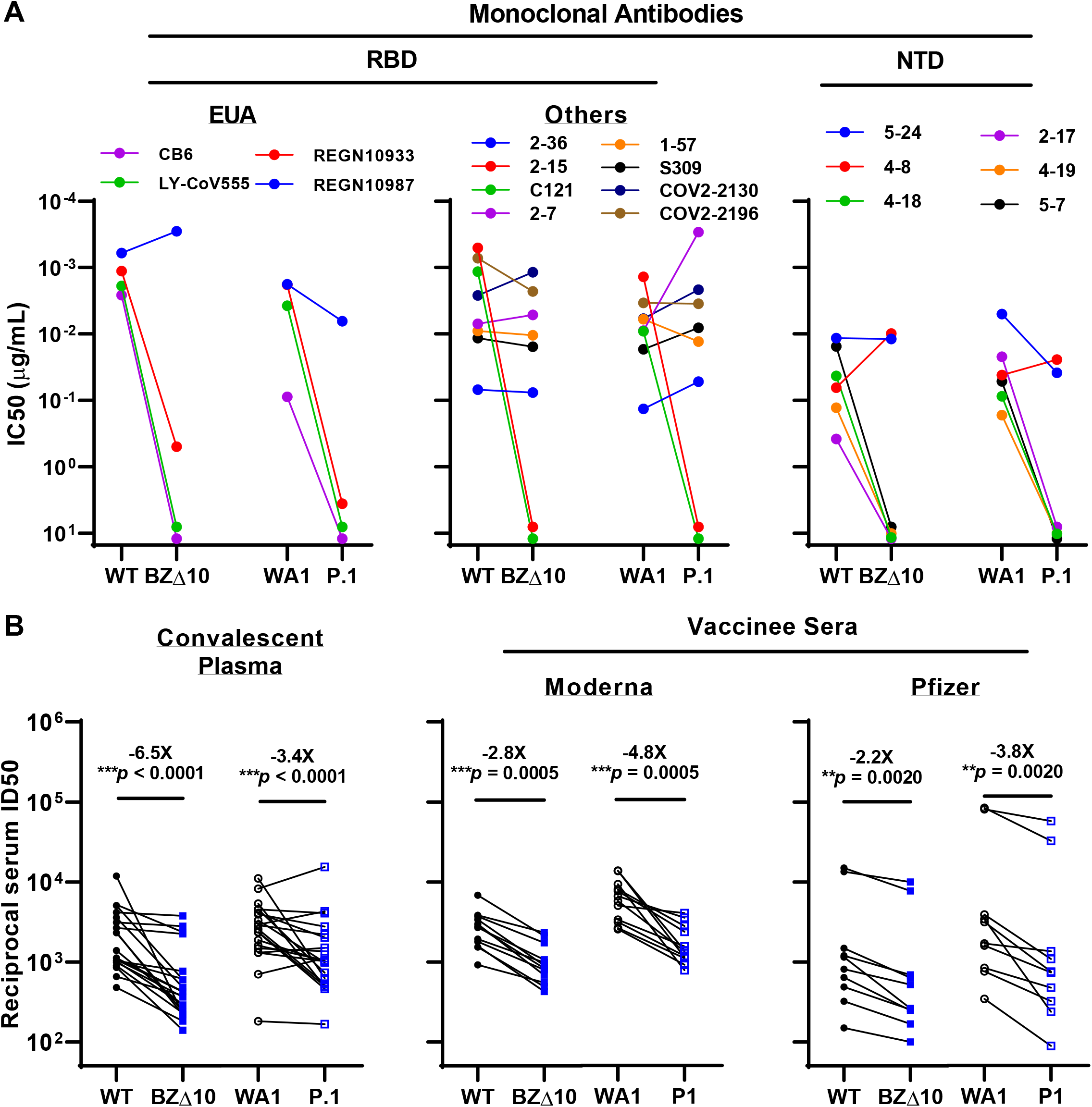
Neutralization of BZΔ10 and P.1 by mAbs, convalescent plasma, and vaccinee sera. See also Figures S1.

a. Changes in neutralization IC50 of select RBD and NTD mAbs.
b. Changes in reciprocal plasma neutralization ID50 values of convalescent plasma and reciprocal serum ID50 values for persons who received Moderna or Pfizer vaccine. Mean fold change in ID50 relative to the WT is written above the *p* values. Statistical analysis was performed using a Wilcoxon matched-pairs signed rank test. Two-tailed p-values are reported.

We also assessed the neutralizing activity of six NTD mAbs (Liu et al., 2020) against the P.1 pseudovirus and authentic virus (Figure 1A, right panel; and Figure S1B). P.1 was profoundly resistant to neutralization by four NTD antibodies: 2-17, 4-18, 4-19, and 5-7. Interestingly, 5-24 and 4-8, two mAbs targeting the antigenic supersite in NTD (Cerutti et al., 2021) that have completely lost neutralizing activity against B.1.351 (Wang et al., 2021), remained active against P.1. To understand the specific mutations responsible for the observed pattern of neutralization, we then tested these NTD mAbs against a panel of pseudoviruses, each containing only a single NTD mutation found in P.1 (Figure S1B). As expected, 5-24 ad 4-8 retained activity against all single-mutation pseudoviruses. P26S only partially accounted for the loss of activity of 4-18; L18F, T20N, and D138Y contributed to the loss of activity of 2-17 and 4-19; and L18F, T20N, D138Y, and R190S together resulted in the loss of activity of 5-7.

We also examined a panel of convalescent plasma obtained from 20 SARS-CoV-2 patients infected in the Spring of 2020, as previously reported (Wang et al., 2021). Each plasma sample was assayed for neutralization against the P.1 pseudovirus and authentic virus in parallel with their WT counterparts. As shown in Figure S1C, many samples lost >2-fold neutralizing activity against BZΔ10 and P.1. The magnitude of the drop in plasma neutralization ID50 titers is summarized in Figure 1B (left panel), showing a 6.5-fold loss of activity against the variant pseudovirus and a 3.4-fold loss of activity against the authentic virus.

Twenty-two vaccinee sera were obtained, as previously reported (Wang et al., 2021), from 12 individuals who received Moderna SARS-Co-2 mRNA-1273 Vaccine (Anderson et al., 2020) and 10 individuals who received the Pfizer BNT162b2 Covid-19 Vaccine (Polack et al., 2020). Each serum sample was assayed for neutralization against BZΔ10 and P.1 together with WT viruses. The extent of the decline in neutralization activity is summarized in Figure 1B (middle and right panels), and each neutralization profile is shown in Figure S1D. A loss of activity against BZΔ10 and P.1 was noted for every sample, but the magnitude of the loss was modest (2.2-2.8 fold for the pseudovirus; 3.8-4.8 fold for the authentic virus) and not as striking as was observed against B.1.351 (6.5-8.6 fold for pseudovirus; 10.3-12.4 fold for authentic virus) (Wang et al., 2021).

To provide insight into the mechanisms of antibody resistance, we determined the structure of the 2-proline-stabilized P.1 spike protein at 3.8 Å resolution by single-particle cryo-EM reconstruction (Figures 2 and S2, Table S1). Overall, the structure of the P.1 spike was highly similar to the D614G variant (Korber et al., 2020; Yurkovetskiy et al., 2020) with 3D classes observed only for the single-RBD-up conformation. This was expected, as the D614G mutation, contained in P1, appears to favor the one-up orientation of RBD, which is required for ACE2 binding and recognition by some RBD-directed antibodies. Structural mobility was observed with the raised RBD (protomer B), but not with protomers A and C, which were in the down orientation (Video S1). Map density was well satisfied by the previously reported single-up structure (PDB 6XM0) for the majority of the trimer, except in three regions. Residues 310-322 in protomer B traced a different path; residues 623-632 were disordered in protomers A and B, and partially ordered in protomer C; and residues 828-853 were disordered in protomers A and C, and partially ordered in protomer B. Notably, two of these regions around residues 320 and 840 were previously observed to “refold” between the single-up and the low-pH all-RBD-down conformation (Zhou et al., 2020), suggesting these regions are generally more mobile – and in this case, sensitive to mutation-induced conformational changes.

**Figure 2.**
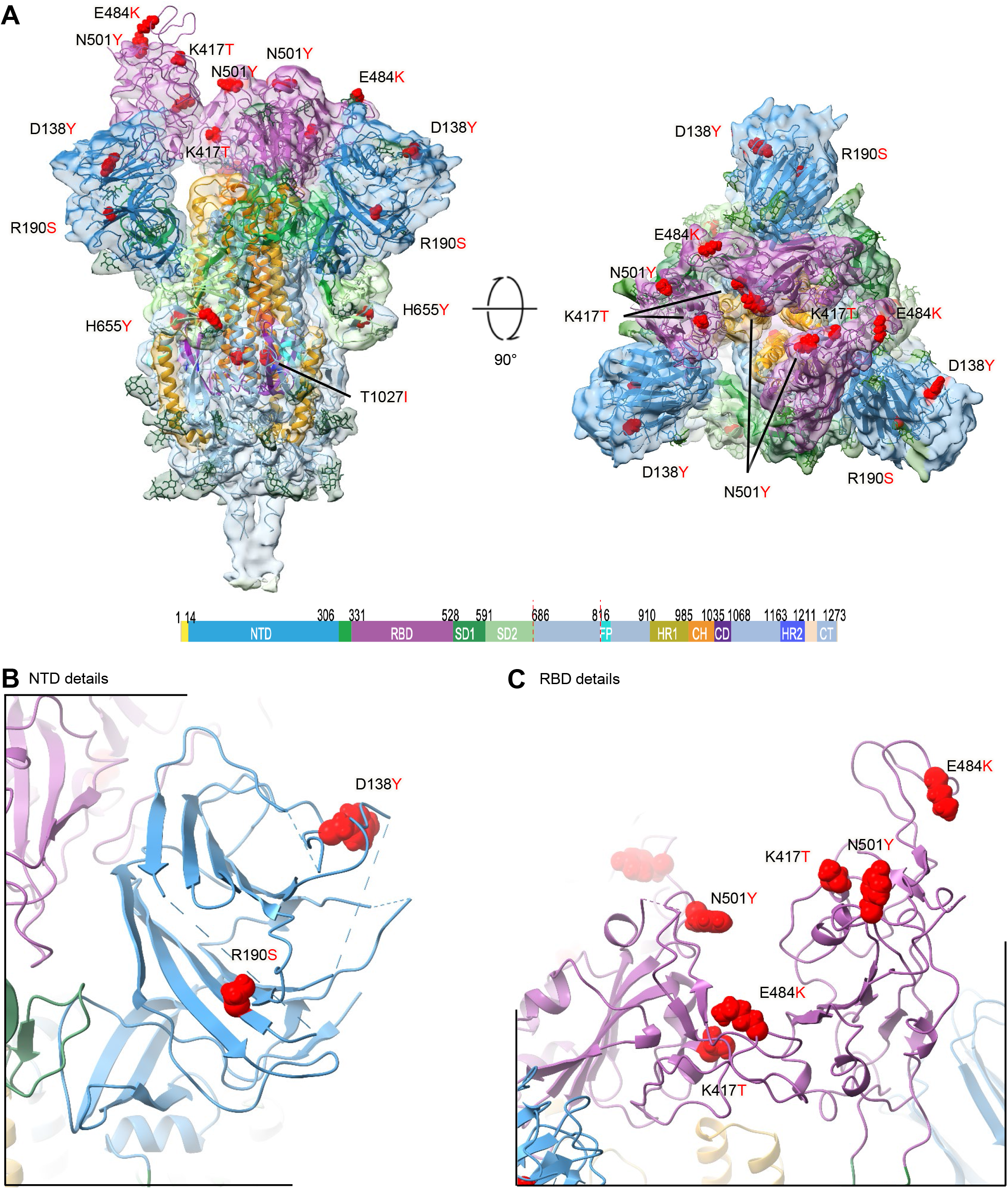
Cryo-EM Structure of the P.1 Spike. See also Figure S2 and Table S1.

a. Overall cryo-EM structure of the P.1 spike trimer with domains colored as shown in key, glycans shown in green, and mutations highlighted in red. Density is shown for the 3.8 Å reconstruction with the molecular model shown in ribbon representation. The left image shows a side view, with viral membrane located below, and the right image shows the view looking down on the spike apex.
b. NTD close up view.
c. RBD close up view.

Because of the high overall conformational similarity to the D614G structure, we infer the functional impact of the P.1 mutation to arise primarily from local changes in structure. Other than H655Y and T1027I, all of the mutations occur within NTD or RBD, which are the targets of neutralizing antibodies. For NTD, the N-terminus was disordered until residue 27, so we were unable to visualize mutations at residue 18, 20, and 26. Mutation D138Y is located in the center of the NTD supersite (Cerutti et al., 2021), explaining its impact on NTD antibodies 2-17 and 4-19 (Fig. S1B), whereas R190S is mostly occluded from the NTD surface (Fig. 2B). For RBD, the three mutations at K417T, E484K and N501Y are all located in the ACE2-binding region and overlap epitopes for multiple neutralizing antibodies. Their relatively equal spatial separation (Fig. 2C) allow them to impact a substantial portion of the ACE2-binding surface.

Overall, the SARS-CoV-2 P.1 variant is of concern because of its rapid rise to dominance as well as its extensive spike mutations, which could lead to antigenic changes detrimental to mAb therapies and vaccine protection. Here we report that P.1 is indeed resistant to neutralization by several RBD-directed mAbs, including three with EUA. The major culprit is the E484K mutation, which has emerged independently in over 50 lineages, including in B.1.526 that we (Annavajhala et al., 2021) and others (West et al., 2021) have identified in New York recently. As for the NTD-directed mAbs, the resistance profiles are markedly different between P.1 and B.1.351, reflecting their distinct sets of mutations in NTD. Both convalescent plasma and vaccinee sera show a significant loss of neutralizing activity against P.1, but the diminution is not as great as that reported against B.1.351 (Garcia-Beltran et al., 2021; Wang et al., 2021). Therefore, the threat of increased re-infection or decreased vaccine protection posed by P.1 may not be as severe as B.1.351. Finally, given that the RBD mutations are largely the same for these two variants, the discrepancy in their neutralization susceptibility to polyclonal plasma or sera suggests that NTD mutations can have a significant effect on the susceptibility of SARS-CoV-2 to antibody neutralization.

## Acknowledgements

The following reagent was obtained through BEI Resources, NIAID, NIH: SARS-Related Coronavirus 2, Isolate hCoV-19/Japan/TY7-503/2021 (Brazil P.1), NR-54982, contributed by National Institute of Infectious Diseases. We thank Bob Grassucci and Chi Wang for help with cryo-EM data collection at the Columbia University cryo-EM Center at the Zuckerman Institute. This study was supported by funding from Andrew & Peggy Cherng, Samuel Yin, Barbara Picower and the JPB Foundation, Brii Biosciences, Roger & David Wu, and the Bill and Melinda Gates Foundation. Support was also provided by the Intramural Program of the Vaccine Research Center, National Institute of Allergy and Infectious Diseases, National Institutes of Health.

## Author contributions

The study was conceptualized by D.D.H. The virology experiments were carried out by P.W., M.S.N., M.W., J.Y., L.L., and Y.H. The structural experiment were carried out by R.G.C., G.C., P.D.K., and L.S. The manuscript was written by P.W., R.G.C., P.D.K., L.S., and D.D.H. and reviewed, commented, and approved by all the authors.

## Competing interests

P.W., J.Y., M.N., Y.H., L.L., and D.D.H. are inventors on a provisional patent application on mAbs to SARS-CoV-2.

## STAR METHODS

## KEY RESOURCES TABLE

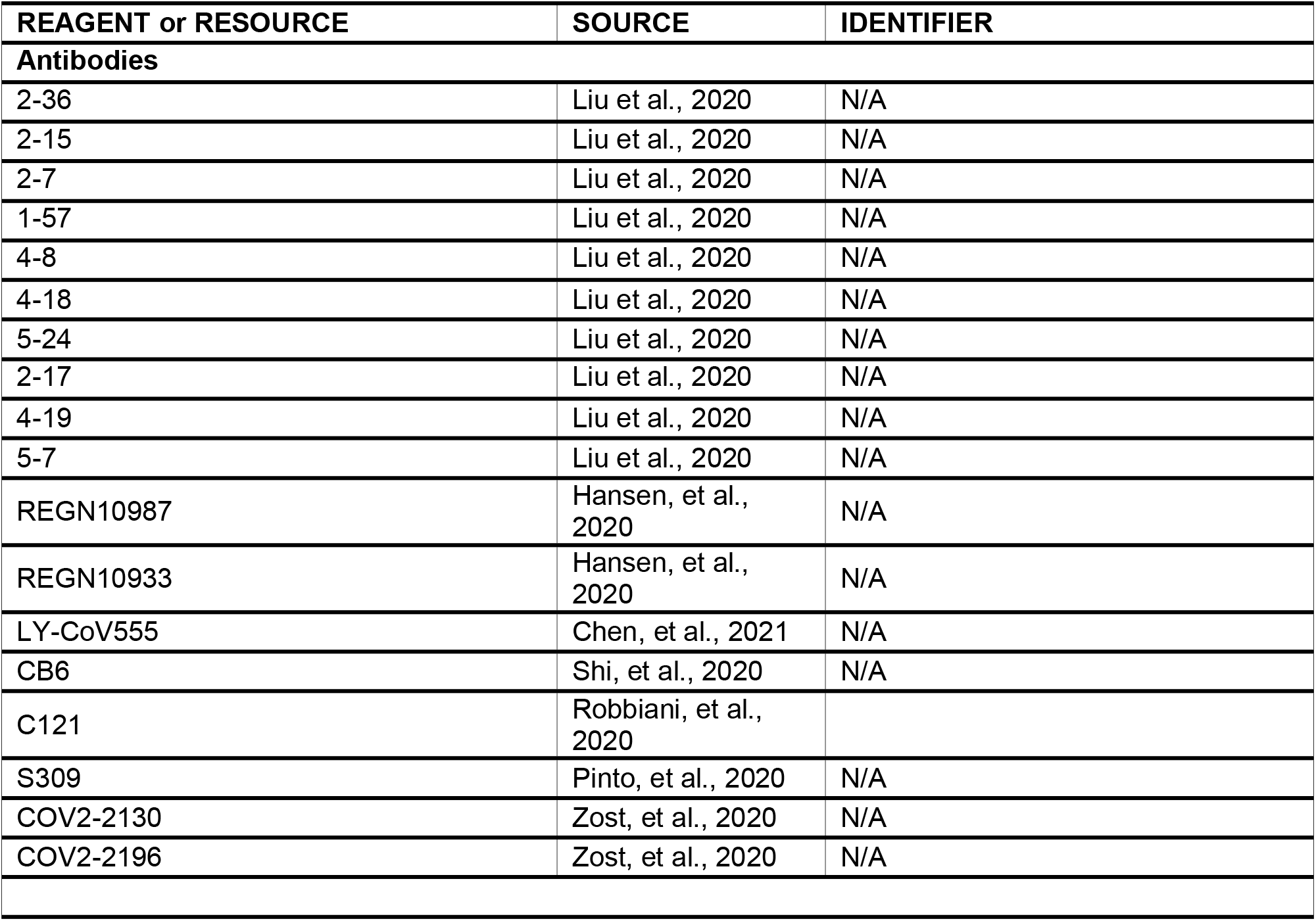
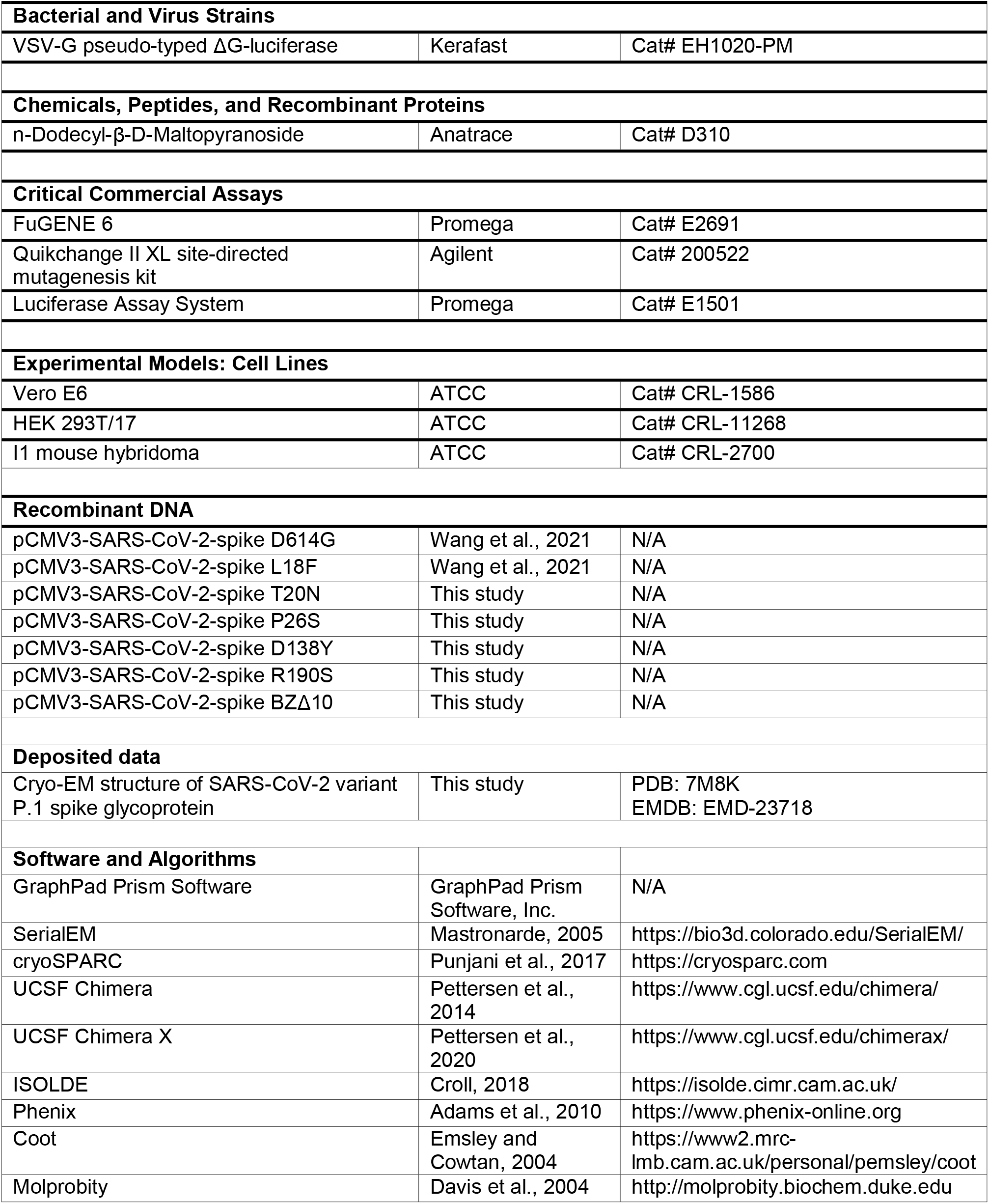

## RESOURCE AVAILABILITY

### Lead contact

Further information and requests for resources and reagents should be directed to and will be fulfilled by the Lead Contact Author David D. Ho.

### Materials availability

All unique/stable reagents generated in this study are available from the Lead Contact with a completed Materials Transfer Agreement.

### Data and code availability

Cryo-EM structure of SARS-CoV-2 variant P.1 spike glycoprotein have been deposited in the PDB (7M8K) and EMDB (EMD-23718).

## EXPERIMENTAL MODEL AND SUBJECT DETAILS

### Cell lines

HEK293T/17 (cat# CRL-11268) and Vero E6 cells (cat# CRL-1586) were from ATCC and cultured in 10% Fetal Bovine Serum (FBS, GIBCO cat# 16140071) supplemented Dulbecco’s Modified Eagle Medium (DMEM, ATCC cat# 30-2002) at 37°C, 5% CO2. I1 mouse hybridoma cells (ATCC, cat# CRL-2700) were cultured in Eagle’s Minimum Essential Medium (EMEM, ATCC cat# 30-2003)) with 20% FBS.

## METHOD DETAILS

### Monoclonal antibodies, patients and vaccinees

Monoclonal antibodies, convalescent plasma, and vaccinee sera were the same as previously reported (Wang et al., 2021).

### Pseudovirus neutralization assays

Plasmids encoding the single-mutation variants found in P.1 and 10-mutation variant (BZΔ10) were generated by Quikchange II XL site-directed mutagenesis kit (Agilent). Recombinant Indiana VSV (rVSV) expressing different SARS-CoV-2 spike variants were generated as previously described (Liu et al., 2020; Wang et al., 2020; Wang et al., 2021). Neutralization assays were performed by incubating pseudoviruses with serial dilutions of mAbs or heat-inactivated plasma or sera, and scored by the reduction in luciferase gene expression as previously described (Liu et al., 2020; Wang et al., 2020; Wang et al., 2021). Briefly, Vero E6 cells (ATCC) were seeded in 96-well plates (2 ×10^4^ cells per well). Pseudoviruses were incubated with serial dilutions of the test samples in triplicate for 30 min at 37□°C. The mixture was added to cultured cells and incubated for an additional 16 hrs. Luminescence was measured using Luciferase Assay System (Promega), and IC_50_ was defined as the dilution at which the relative light units were reduced by 50% compared with the virus control wells (virus + cells) after subtraction of the background in the control groups with cells only. The IC_50_ values were calculated using a five-parameter dose-response curve in GraphPad Prism.

### Authentic SARS-CoV-2 Microplate Neutralization

The SARS-CoV-2 viruses USA-WA1/2020 (WA1), and hCoV-19/Japan/TY7-503/2021 (P.1) were obtained from BEI Resources (NIAID, NIH). The deposited virus (Passage 2 in Vero E6/TMPRSS2 cells) was reported to have an additional mutation as compared to the clinical isolate: NSP6 (Non-structural protein 6) F184V (GISAID: EPI_ISL_877769). The viruses were propagated for one passage using Vero E6 cells. Virus infectious titer was determined by an end-point dilution and cytopathic effect (CPE) assay on Vero E6 cells as described previously (Liu et al., 2020; Wang et al., 2020; Wang et al., 2021).

An end-point-dilution microplate neutralization assay was performed to measure the neutralization activity of convalescent plasma samples, vaccinee sera, and purified mAbs. Triplicates of each dilution were incubated with SARS-CoV-2 at an MOI of 0.1 in EMEM with 7.5% inactivated fetal calf serum (FCS) for 1 hour at 37°C. Post incubation, the virus-antibody mixture was transferred onto a monolayer of Vero E6 cells grown overnight. The cells were incubated with the mixture for ∼70 hrs. CPE was visually scored for each well in a blinded fashion by two independent observers. The results were then converted into percentage neutralization at a given sample dilution or mAb concentration, and the averages ± SEM were plotted using a five-parameter dose-response curve in GraphPad Prism.

### Cryo-EM data collection and processing

2µL P.1 spike protein at a concentration of 2 mg/ml was incubated in 10 mM Hepes pH 7.4, 150 mM NaCl, and 0.005% n-dodecyl-β-D-maltoside (DDM) was incubated on C-flat 1.2/1.3 carbon grids for 30 seconds and vitrified using a Vitrobot plunge freezer. Data were collected on a Titan Krios electron microscope operating at 300 kV, equipped with a Gatan K3 direct electron detector and energy filter, using the Leginon software package (Suloway et al., 2005). A total electron fluence of 51.69 e-/Å2 was fractionated over 40 frames, with a total exposure time of 2.0 seconds. A magnification of 81,000x resulted in a pixel size of 1.058 Å, and a defocus range of -0.4 to -3.5 µm was used.

All processing was done using cryoSPARC v2.15.0 (Punjani et al., 2017). Raw movies were aligned and dose-weighted using patch motion correction, and the CTF was estimated using patch CTF estimation. A small subset of approximately 200 micrographs were picked using blob picker, followed by 2D classification and manual curation of particle picks, and used to train a Topaz neural network. This network was then used to pick particles from the remaining micrographs, which were extracted with a box size of 384 pixels.

### Cryo-EM model building

We used PDB 6XM0, one of the most complete coronavirus spike structures, as a starting model. The model was docked to the map using Chimera. The model was then fitted interactively using COOT (Emsley and Cowtan, 2004) with real space refinement performed in Phenix 1.18 (Adams et al., 2004). Validation was performed using Molprobity (Davis et al., 2004) and EMRinger (Barad et al., 2015). The model was submitted to the PDB with PDB ID XXX. Figures were prepared using UCSF ChimeraX (Goddard et al., 2018).

**Figure S1. Neutralization of BZ**Δ**10 and P.1 by mAbs, convalescent plasma, and vaccinee sera**. Related to Figure 1

a. Neutralization of WT and BZ_△_10 pseudoviruses, and WA1 and P.1 authentic viruses by anti-RBD mAbs.
b. Neutralization of WT, BZ_△_10, and single-mutation pseudoviruses, and WA1 and P.1 authentic viruses by anti-NTD mAbs.
c. Neutralization of WT and BZ_△_10 pseudoviruses, and WA1 and P.1 authentic viruses by convalescent plasma.
d. Neutralization of WT and BZ_△_10 pseudoviruses, and WA1 and P.1 authentic viruses by vaccinee sera. Data represent mean ± SEM of technical triplicates.

**Figure S2. Cryo-EM Data Collection and Refinement**. Related to Figure 2

a. Representative micrograph, power spectrum, and contrast transfer function (CTF) fit.
b. Representative 2D class averages showing spike particles.
c. Global consensus refinement Fourier Shell Correlation (FSC) curve and particle projection viewing angle distribution.
d. Local resolution estimation mapped on surface density for global refinement.

**Table S1. Cryo-EM Data Collection and Refinement Statistics**. Related to Figure 2

**Video S1. Cryosparc 3D-variability analysis of the P1 spike**. Related to Figure 2

## Notes

### Competing Interest Statement

P.W., J.Y., M.N., Y.H., and D.D.H. are inventors on a provisional patent application on mAbs to SARS-CoV-2.

### Summary of Updates

The Cryo-EM structure of P.1 spike trimer added.

## References

Adams, P.D., Gopal, K., Grosse-Kunstleve, R.W., Hung, L.W., Ioerger, T.R., McCoy, A.J., Moriarty, N.W., Pai, R.K., Read, R.J., Romo, T.D., et al. (2004). Recent developments in the PHENIX software for automated crystallographic structure determination. J Synchrotron Radiat 11, 53–55.

Anderson, E.J., Rouphael, N.G., Widge, A.T., Jackson, L.A., Roberts, P.C., Makhene, M., Chappell, J.D., Denison, M.R., Stevens, L.J., Pruijssers, A.J., et al. (2020). Safety and Immunogenicity of SARS-CoV-2 mRNA-1273 Vaccine in Older Adults. N Engl J Med 383, 2427–2438.

Annavajhala, M.K., Mohri, H., Zucker, J.E., Sheng, Z., Wang, P., Gomez-Simmonds, A., Ho, D.D., and Uhlemann, A.-C. (2021). A Novel SARS-CoV-2 Variant of Concern, B.1.526, Identified in New York. Preprint at https://www.medrxiv.org/content/medrxiv/early/2021/2002/2025/2021.2002.2023.21252259.

Barad, B.A., Echols, N., Wang, R.Y., Cheng, Y., DiMaio, F., Adams, P.D., and Fraser, J.S. (2015). EMRinger: side chain-directed model and map validation for 3D cryo-electron microscopy. Nat Methods 12, 943–946.

Cerutti, G., Guo, Y., Zhou, T., Gorman, J., Lee, M., Rapp, M., Reddem, E.R., Yu, J., Bahna, F., Bimela, J., et al. (2021). Potent SARS-CoV-2 Neutralizing Antibodies Directed Against Spike N-Terminal Domain Target a Single Supersite. Preprint at https://www.biorxiv.org/content/biorxiv/early/2021/2001/2011/2021.2001.2010.426120.

Chen, P., Nirula, A., Heller, B., Gottlieb, R.L., Boscia, J., Morris, J., Huhn, G., Cardona, J., Mocherla, B., Stosor, V., et al. (2021). SARS-CoV-2 Neutralizing Antibody LY-CoV555 in Outpatients with Covid-19. N Engl J Med 384, 229–237.

Davis, I.W., Murray, L.W., Richardson, J.S., and Richardson, D.C. (2004). MOLPROBITY: structure validation and all-atom contact analysis for nucleic acids and their complexes. Nucleic Acids Res 32, W615–619.

Emsley, P., and Cowtan, K. (2004). Coot: model-building tools for molecular graphics. Acta Crystallogr D Biol Crystallogr 60, 2126–2132.

Faria, N.R. (2021). Genomic characterisation of an emergent SARS-CoV-2 lineage in Manaus: preliminary findings. https://virological.org/t/genomic-characterisation-of-an-emergent-sars-cov-2-lineage-in-manaus-preliminary-findings/586

Garcia-Beltran, W.F., Lam, E.C., Denis, K.S., Nitido, A.D., Garcia, Z.H., Hauser, B.M., Feldman, J., Pavlovic, M.N., Gregory, D.J., Poznansky, M.C., et al. (2021). Circulating SARS-CoV-2 variants escape neutralization by vaccine-induced humoral immunity. Preprint at https://www.medrxiv.org/content/medrxiv/early/2021/2002/2018/2021.2002.2014.21251704.

Goddard, T.D., Huang, C.C., Meng, E.C., Pettersen, E.F., Couch, G.S., Morris, J.H., and Ferrin, T.E. (2018). UCSF ChimeraX: Meeting modern challenges in visualization and analysis. Protein Sci 27, 14–25.

Gottlieb, R.L., Nirula, A., Chen, P., Boscia, J., Heller, B., Morris, J., Huhn, G., Cardona, J., Mocherla, B., Stosor, V., et al. (2021). Effect of Bamlanivimab as Monotherapy or in Combination With Etesevimab on Viral Load in Patients With Mild to Moderate COVID-19: A Randomized Clinical Trial. JAMA 325, 632–644.

Hansen, J., Baum, A., Pascal, K.E., Russo, V., Giordano, S., Wloga, E., Fulton, B.O., Yan, Y., Koon, K., Patel, K., et al. (2020). Studies in humanized mice and convalescent humans yield a SARS-CoV-2 antibody cocktail. Science 369, 1010–1014.

Korber, B., Fischer, W.M., Gnanakaran, S., Yoon, H., Theiler, J., Abfalterer, W., Hengartner, N., Giorgi, E.E., Bhattacharya, T., Foley, B., et al. (2020). Tracking Changes in SARS-CoV-2 Spike: Evidence that D614G Increases Infectivity of the COVID-19 Virus. Cell 182, 812–827 e819.

Liu, L., Wang, P., Nair, M.S., Yu, J., Rapp, M., Wang, Q., Luo, Y., Chan, J.F., Sahi, V., Figueroa, A., et al. (2020). Potent neutralizing antibodies against multiple epitopes on SARS-CoV-2 spike. Nature 584, 450–456.

Liu, Y., Liu, J., Xia, H., Zhang, X., Fontes-Garfias, C.R., Swanson, K.A., Cai, H., Sarkar, R., Chen, W., Cutler, M., et al. (2021). Neutralizing Activity of BNT162b2-Elicited Serum - Preliminary Report. N Engl J Med, doi: 10.1056/NEJMc2102017. Online ahead of print.

Naveca, F. (2021). Phylogenetic relationship of SARS-CoV-2 sequences from Amazonas with emerging Brazilian variants harboring mutations E484K and N501Y in the Spike protein. https://virological.org/t/phylogenetic-relationship-of-sars-cov-2-sequences-from-amazonas-with-emerging-brazilian-variants-harboring-mutations-e484k-and-n501y-in-the-spike-protein/585.

Pinto, D., Park, Y.-J., Beltramello, M., Walls, A.C., Tortorici, M.A., Bianchi, S., Jaconi, S., Culap, K., Zatta, F., De Marco, A., et al. (2020). Cross-neutralization of SARS-CoV-2 by a human monoclonal SARS-CoV antibody. Nature 583, 290–295.

Polack, F.P., Thomas, S.J., Kitchin, N., Absalon, J., Gurtman, A., Lockhart, S., Perez, J.L., Perez Marc, G., Moreira, E.D., Zerbini, C., et al. (2020). Safety and Efficacy of the BNT162b2 mRNA Covid-19 Vaccine. N Engl J Med 383, 2603–2615.

Punjani, A., Rubinstein, J.L., Fleet, D.J., and Brubaker, M.A. (2017). cryoSPARC: algorithms for rapid unsupervised cryo-EM structure determination. Nat Methods 14, 290–296.

Robbiani, D.F., Gaebler, C., Muecksch, F., Lorenzi, J.C.C., Wang, Z., Cho, A., Agudelo, M., Barnes, C.O., Gazumyan, A., Finkin, S., et al. (2020). Convergent antibody responses to SARS-CoV-2 in convalescent individuals. Nature 584, 437–442.

Shi, R., Shan, C., Duan, X., Chen, Z., Liu, P., Song, J., Song, T., Bi, X., Han, C., Wu, L., et al. (2020). A human neutralizing antibody targets the receptor-binding site of SARS-CoV-2. Nature 584, 120–124.

Suloway, C., Pulokas, J., Fellmann, D., Cheng, A., Guerra, F., Quispe, J., Stagg, S., Potter, C.S., and Carragher, B. (2005). Automated molecular microscopy: the new Leginon system. J Struct Biol 151, 41–60.

Tegally, H., Wilkinson, E., Giovanetti, M., Iranzadeh, A., Fonseca, V., Giandhari, J., Doolabh, D., Pillay, S., San, E.J., Msomi, N., et al. (2021). Emergence of a SARS-CoV-2 variant of concern with mutations in spike glycoprotein. Nature.

Wang, P., Liu, L., Nair, M.S., Yin, M.T., Luo, Y., Wang, Q., Yuan, T., Mori, K., Solis, A.G., Yamashita, M., et al. (2020). SARS-CoV-2 neutralizing antibody responses are more robust in patients with severe disease. Emerg Microbes Infect 9, 2091–2093.

Wang, P., Nair, M.S., Liu, L., Iketani, S., Luo, Y., Guo, Y., Wang, M., Yu, J., Zhang, B., Kwong, P.D., et al. (2021). Antibody Resistance of SARS-CoV-2 Variants B.1.351 and B.1.1.7. Nature, In press.

West, A.P., Barnes, C.O., Yang, Z., and Bjorkman, P.J. (2021). SARS-CoV-2 lineage B.1.526 emerging in the New York region detected by software utility created to query the spike mutational landscape. Preprint at https://www.biorxiv.org/content/biorxiv/early/2021/2002/2023/2021.2002.2014.431043.

Wu, K., Werner, A.P., Koch, M., Choi, A., Narayanan, E., Stewart-Jones, G.B.E., Colpitts, T., Bennett, H., Boyoglu-Barnum, S., Shi, W., et al. (2021). Serum Neutralizing Activity Elicited by mRNA-1273 Vaccine - Preliminary Report. N Engl J Med, doi: 10.1056/NEJMc2102179. Online ahead of print.

Yurkovetskiy, L., Wang, X., Pascal, K.E., Tomkins-Tinch, C., Nyalile, T.P., Wang, Y., Baum, A., Diehl, W.E., Dauphin, A., Carbone, C., et al. (2020). Structural and Functional Analysis of the D614G SARS-CoV-2 Spike Protein Variant. Cell 183, 739–751 e738.

Zhou, T., Tsybovsky, Y., Gorman, J., Rapp, M., Cerutti, G., Chuang, G.Y., Katsamba, P.S., Sampson, J.M., Schon, A., Bimela, J., et al. (2020). Cryo-EM Structures of SARS-CoV-2 Spike without and with ACE2 Reveal a pH-Dependent Switch to Mediate Endosomal Positioning of Receptor-Binding Domains. Cell Host Microbe 28, 867–879 e865.

Zost, S.J., Gilchuk, P., Case, J.B., Binshtein, E., Chen, R.E., Nkolola, J.P., Schafer, A., Reidy, J.X., Trivette, A., Nargi, R.S., et al. (2020). Potently neutralizing and protective human antibodies against SARS-CoV-2. Nature 584, 443–449.

