## Supplemental text and figures for "Increased Resistance of SARS-CoV-2 Variant P.1 to Antibody Neutralization"

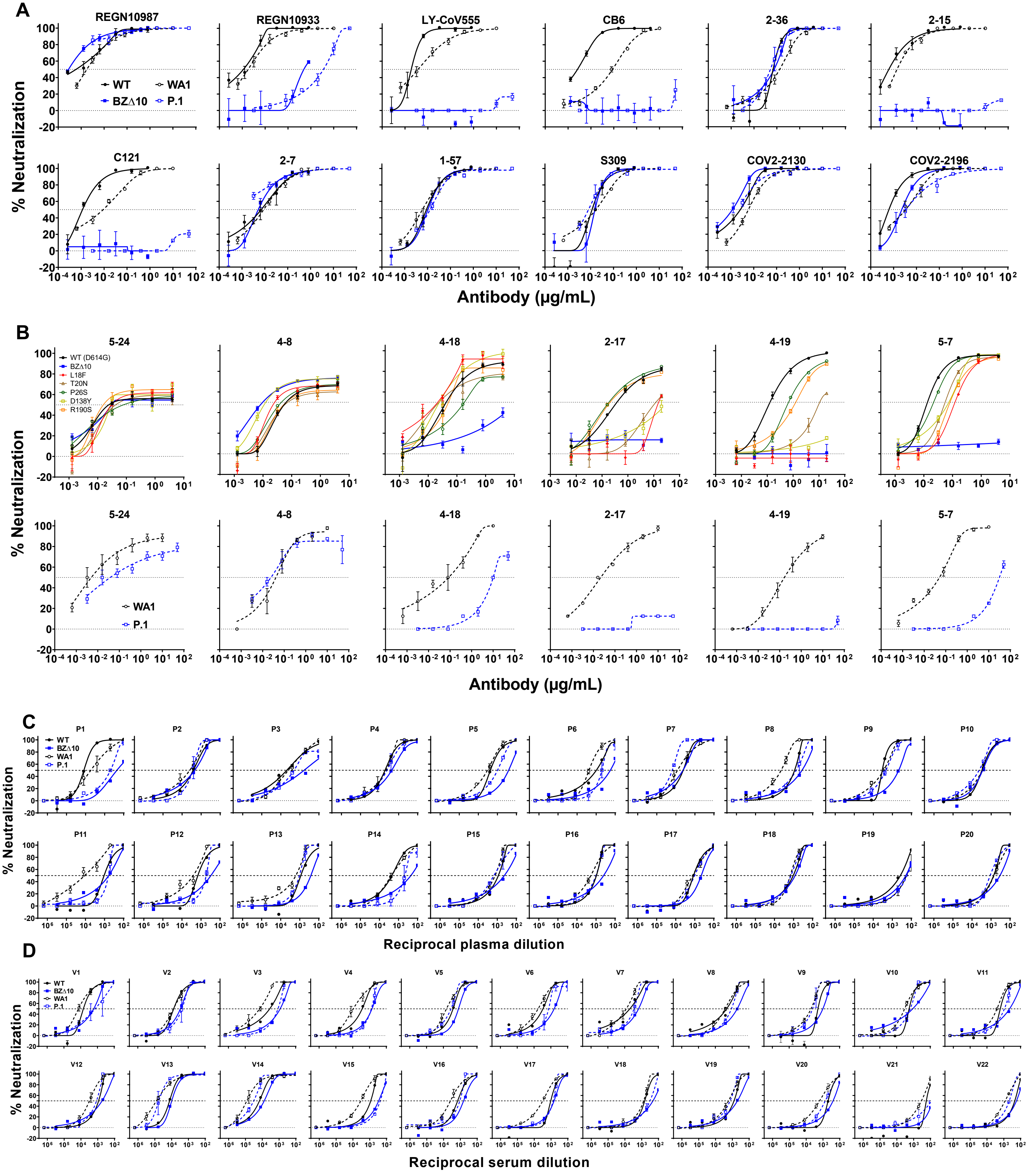

A
REGN10987
REGN10933
LY-CoV555
CB6
2-36
2-15
C121
2-7
1-57
S309
COV2-2130
COV2-2196
WT
WA1
BZΔ10
P.1
% Neutralization
Antibody (μg/mL)
B
5-24
4-8
4-18
2-17
4-19
5-7
WT (D614G)
BZΔ10
L18F
T20N
P26S
D138Y
R190S
WA1
P.1
% Neutralization
Antibody (μg/mL)
C
P1
P2
P3
P4
P5
P6
P7
P8
P9
P10
P11
P12
P13
P14
P15
P16
P17
P18
P19
P20
% Neutralization
Reciprocal plasma dilution
D
V1
V2
V3
V4
V5
V6
V7
V8
V9
V10
V11
V12
V13
V14
V15
V16
V17
V18
V19
V20
V21
V22
% Neutralization
Reciprocal serum dilution

**Figure S1. Neutralization of BZΔ10 and P.1 by mAbs, convalescent plasma, and vaccinee sera.** related to Figure 1
(A) Neutralization of WT and BZ△10 pseudoviruses, and WA1 and P.1 authentic viruses by anti-RBD mAbs.
(B) Neutralization of WT, BZ△10, and single-mutation pseudoviruses, and WA1 and P.1 authentic viruses by anti-NTD mAbs.
(C) Neutralization of WT and BZ△10 pseudoviruses, and WA1 and P.1 authentic viruses by convalescent plasma.
(D) Neutralization of WT and BZ△10 pseudoviruses, and WA1 and P.1 authentic viruses by vaccinee sera.
Data represent mean ± SEM of technical triplicates.

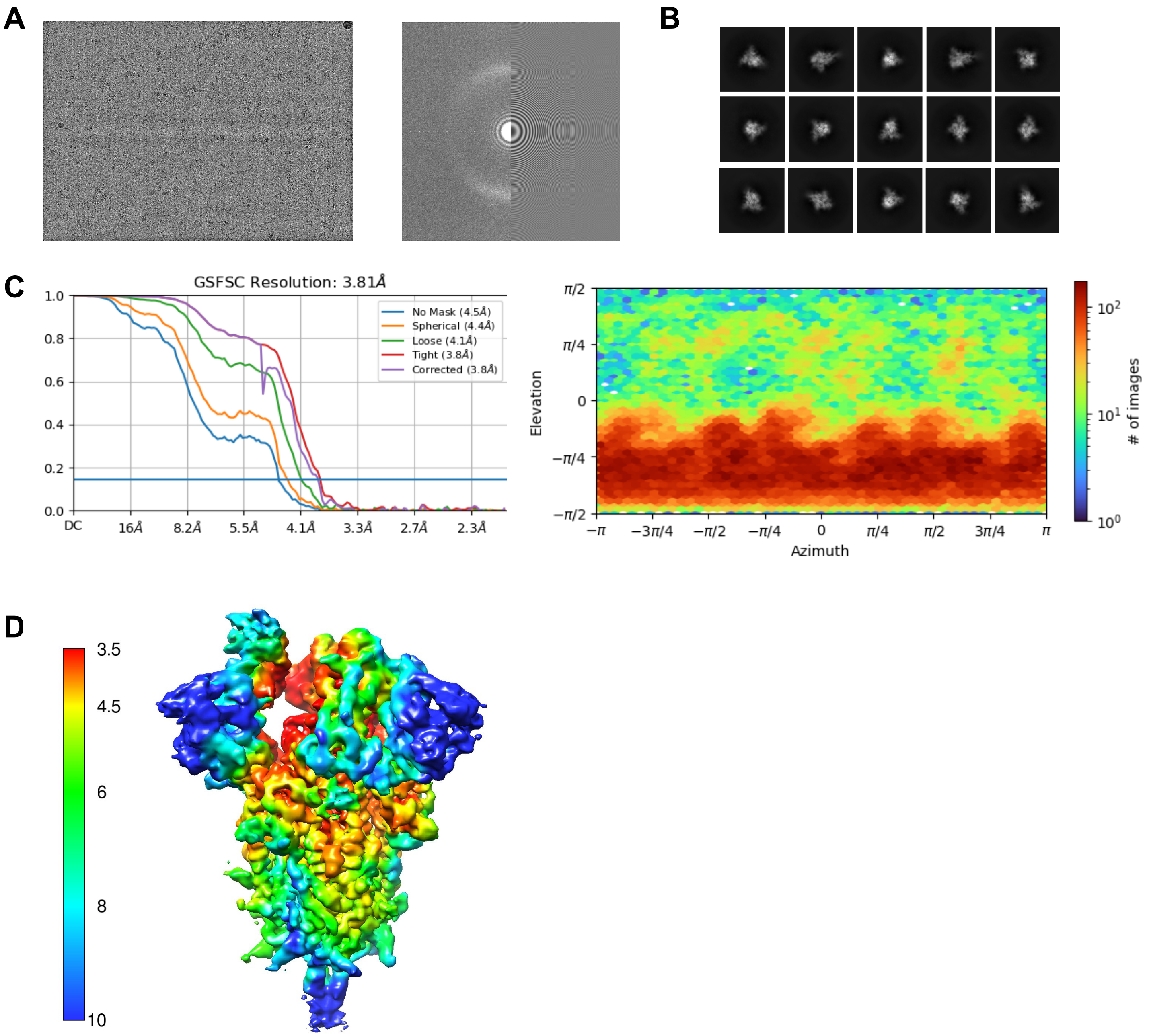


**Figure S2. Cryo-EM Data Collection and Refinement.** related to Figure 2

(A) Representative micrograph, power spectrum, and contrast transfer function (CTF) fit.

(B) Representative 2D class averages showing spike particles.

(C) Global consensus refinement Fourier Shell Correlation (FSC) curve and particle projection viewing angle distribution.

(D) Local resolution estimation mapped on surface density for global refinement.

**Table S1. Cryo-EM Data Collection and Refinement Statistics (Related to Figure 2)**

| | SARS-CoV-2 variant P.1 spike |
|---|---|
| **EMDB ID** | EMD-23718 |
| **PDB ID** | 7M8K |
| Data Collection | |
| Microscope | FEI Titan Krios |
| Voltage (kV) | 300 |
| Electron dose ($e^-/Å^2$) | 48 |
| Detector | Gatan K3 BioQuantum |
| Pixel Size (Å) | 1.07 |
| Defocus Range (µm) | -0.8/-2.5 |
| Magnification | 81000 |
| Reconstruction | |
| Software | cryoSPARC v3.1 |
| Particles | 120,543 |
| Symmetry | C1 |
| Box size (pix) | 384 |
| Resolution (Å) ($FSC_{0.143}$) | 3.81 |
| Refinement | |
| Software | Phenix 1.18 |
| Protein residues | 3084 |
| Chimera CC | 0.82 |
| R.m.s. deviations | |
| Bond lengths (Å) | 0.008 |
| Bond angles (°) | 1.523 |
| Validation | |
| Molprobity score | 1.3 |
| Clash score | 3.10 |
| Favored rotamers (%) | 99.89 |
| Ramachandran | |
| Favored regions (%) | 96.8 |
| Allowed regions (%) | 3.2 |
| Disallowed regions (%) | 0 |
